# Structural determinants of signal speed: Estimated axonal latency and its multimodal validation during face processing in autism

**DOI:** 10.1101/2025.03.19.644214

**Authors:** Campbell R. Coleman, Madelyn G. Nance, Zachary Jacokes, T. Jason Druzgal, Vardan Arutiunian, Anna Kresse, Catherine A.W. Sullivan, Megha Santhosh, Emily Neuhaus, Heather Borland, Raphael A. Bernier, Susan Y. Bookheimer, Mirella Dapretto, Allison Jack, Natalia M. Kleinhans, Shafali Jeste, James C. McPartland, Adam Naples, Daniel Geschwind, Abha R. Gupta, Sara Jane Webb, Kevin A. Pelphrey, John Darrell Van Horn, Benjamin T. Newman, Meghan H. Puglia, ACE GENDAAR Consortium

## Abstract

It has not previously been possible to investigate the fundamental relationship between axonal structure – which dictates action potential transmission – and human neuronal function *in vivo*. Here, we introduce a novel metric of axonal signal speed, estimated axonal latency (EAL), derived from the relationship between axonal diameter, myelination, and length measured via MRI. We validate EAL along two pathways of the face processing network by relating it to N170 latency, an electrophysiological marker of face processing speed measured via EEG. Our results show that EAL along these pathways predicts N170 latency specifically during face processing. Moreover, we demonstrate that individuals with and without autism rely upon different pathways, potentially providing a structural account for autism-related face processing differences. By establishing this relationship between EEG-based electrical function and MRI-based axonal microstructure, we provide a non-invasive, spatially detailed estimate of neuronal processing speed that can inform our understanding of brain function, development, and disorder.

**Teaser:** Estimated axonal latency is a non-invasive, spatially detailed measure of neuronal speed to inform brain function and disorder.

## Introduction

The fundamental method for neuronal communication is action potential transmission via the axon(*1*). This process is intrinsic to brain function at the cellular, circuit, and system level. Action potential transmission speed is crucial for network dynamics as the brain must rapidly integrate input from complex stimuli to generate cognition and behavior. The primary constraints on action potential transmission arise from the structural composition of the axon. Insulating myelin and increased fiber size reduce external and internal electrical resistance to enable faster transmission, communication, and processing speed(*2*, *3*). These processes are essential to successful neurodevelopment(*4*) and are often altered in neurological disorders(*5*).

Although animal studies of action potential generation in peripheral nerves have clearly demonstrated the fundamental relationship between axonal structure and neuronal function(*6*, *7*), this relationship has not yet been investigated in the human central nervous system. Here, we apply multimodal neuroimaging in adolescents with and without autism to relate microstructural differences in white matter measured via diffusion magnetic resonance imaging (dMRI) to electrophysiological signals underlying cognitive processes measured via electroencephalography (EEG). By establishing this relationship between EEG-based neuronal electrical function and MRI-based axonal microstructure, this study provides a non-invasive, spatially-detailed means to predict action potential speed between different areas of the brain, opening the door to regionally and individually-specific estimates of neuronal processing speed.

EEG is a useful tool for studying cognitive processes with high temporal resolution. The event-related potential (ERP) technique is one of the most common analytic methods applied to EEG data. ERPs are stereotyped neural responses evoked via sensory stimuli and/or cognitive processes(*8*). Researchers can use the ERP technique to compute the latency, amplitude, and waveform of a signal, but cannot visualize the structural integrity of the ensemble of neurons that created the EEG signal(*9*). Conversely, MRI is a modality with exceptional spatial resolution that provides detailed information about brain structure and connectivity, but only provides an indirect and relatively slow measure of brain function via the proxy of blood oxygenation(*10*). Multimodal neuroimaging techniques combining EEG and MRI allow us to capitalize on both the temporal and spatial strengths of each technique(*11*). Here, using a well established ERP metric, we validate a novel diffusion imaging estimate of electrical function – estimated axonal latency (EAL) – to establish the relationship between axonal structure and signal speed in the brain.

G-ratio is a voxel-wise measure of axonal geometry derived from dMRI that is proportional to the ratio of inner axonal diameter to total axonal diameter(*12*). This measure encapsulates the competing contributions of myelin thickness and inner axonal diameter, which together serve to insulate the axon, reduce electrical resistance, and allow for faster action potential transmission(*13*). G-ratio can be mathematically transformed to a second metric called aggregate conduction velocity(*14*), an indirect, large-scale approximation of voxel-wise neuronal action potential speed in cortical white matter. To validate our ability to measure *in vivo* signal speed transmission without the use of invasive electrophysiological techniques, we introduce EAL, a metric derived from the physical relationship between speed (i.e., aggregate conduction velocity) and distance (i.e., white matter tract length between two regions of the brain). By applying this novel metric to adolescent populations with and without autism, we demonstrate the ability of EAL to account for differences in neuronal processing speed that occur in development and vary with disorder.

To validate EAL as a metric of neuronal signal speed, we compare its association to electrophysiological latency during face processing. We chose to focus on face processing as it is a well-defined cognitive process with known neurological correlates and important implications for human evolution(*15*) and behavior(*16*). The N170 is a stereotyped electrophysiological signal evoked approximately 170 milliseconds after the presentation of a visual stimulus that typically shows greater negativity to faces(*17*–*19*). N170 latency decreases with development(*20*) and differs in autism spectrum disorder (ASD)(*21*–*25*), a highly heterogeneous neurodevelopmental disorder characterized by differences in brain structure and cognitive function(*26*) including face processing(*27*). Some researchers have attributed N170 latency differences in ASD to atypical patterns of visual attention(*28*) or generally slower visual processing(*29*). However, it is important to consider that these N170 latency differences are inconsistent(*30*–*33*), and each of these explanations fails to consider underlying neural structure.

Here, we investigate the relationship between N170 latency and EAL along two white matter pathways commonly associated with face processing(*34*, *35*) and putatively implicated in N170 signal generation via the ventral visual processing stream(*21*, *36*, *37*): the primary visual cortex to right fusiform gyrus (V1-rFG) and the primary visual cortex to right posterior superior temporal sulcus (V1-rpSTS). By linking EAL to an established electrophysiological biomarker of face processing, this study provides a critical bridge between structural and functional aspects of neural processing. Our findings advance our understanding of how white matter microstructure contributes to cognitive function and offer a promising framework for identifying biologically grounded, individualized markers of processing time.

## Results

As part of a multisite National Institute of Health sponsored Autism Center for Excellence Network, 125 adolescents (Table 1) underwent comprehensive phenotyping, EEG, and structural and diffusion MRI. While undergoing EEG, participants were presented with faces and symbols as feedback on their performance during an implicit learning task(*38*). After preprocessing the EEG data(*39*), we computed N170 latency as the negative peak latency occurring 170-270 ms after stimulus onset (Fig. 1A) in posterior/occipital channels over the right hemisphere (Fig. 1B) for each condition (faces, symbols). Participants also underwent diffusion, T1-weighted, and T2-weighted imaging. Resulting images were processed per the protocol described in Newman et al.(*40*). We defined V1, rFG, and rpSTS regions of interest (Fig. 1C) using the term-based meta-analytic approach available via NeuroSynth.org, and investigated tractograms along the V1-rFG and V1-rpSTS pathways (Fig. 1D). We then generated aggregate g-ratio and aggregate conduction velocity maps for each of our pathways of interest (Fig. 1E), and computed EAL for each pathway by dividing aggregate conduction velocity by tract length.

**Fig. 1:**
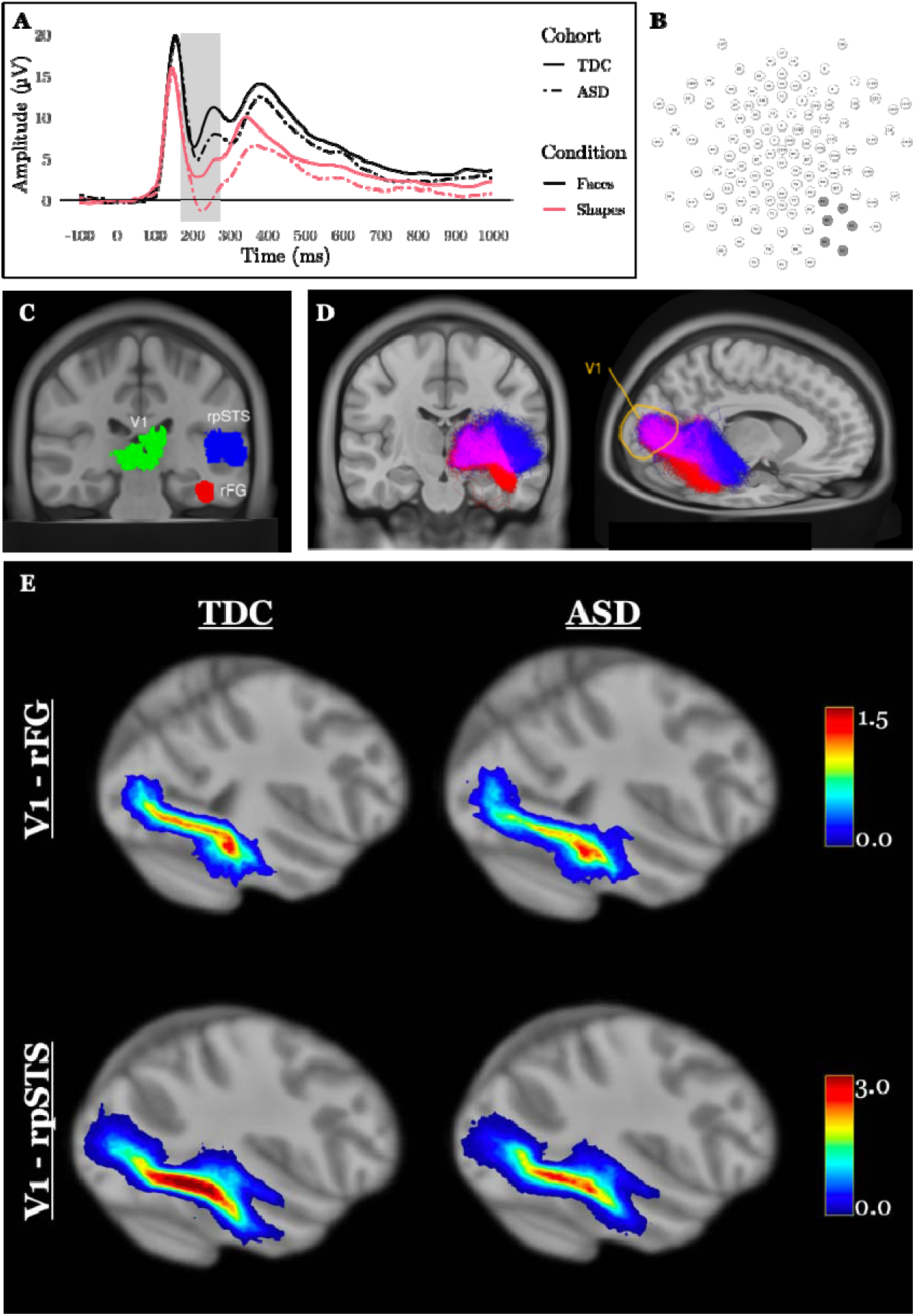
A multimodal investigation of the relationship between electrical function and axonal structure during face processing. **A)** To measure EEG-based electrical function during face processing, TDC (solid line) and ASD (dashed line) participants underwent EEG during the perception of faces (black) and shapes (pink). We computed N170 latency as the negative peak latency 170-270 ms after stimulus onset (grey shading). **B)** We extracted N170 latency from right posterior/occipital (shaded) channels. **C)** To extract MRI-based axonal microstructure, we generated regions of interest including primary visual cortex (V1, green), right fusiform gyrus (rFG, red), and right posterior superior temporal sulcus (rpSTS, blue) using a meta-analytic approach. **D)** Axonal microstructure was measured along two pathways, V1-rFG (red) and V1-rpSTS (blue). Overlapping tracts are pictured in magenta. **E)** Mean aggregate conduction velocity maps along all tracts retained in the study are separated by diagnostic group and tract. To create each map, voxels traversed by a retained tract were summed across all participants in a group, then divided by the total number of participants for a representation of within-group tract cohesiveness and overall conduction velocity.

**Table 1:** Participant Demographics. Demographic information was recorded at each site. Total brain volume measurements were obtained from T1-weighted images. There are no significant group differences across TDC and ASD cohorts for any of our considered demographic variables.

| Demographic |  | Total Sample | TDC Cohort | ASD Cohort |
| --- | --- | --- | --- | --- |
| Age (Years) | Range | 8.08 - 17.92 | 8.08-17.92 | 8.17 - 17.83 |
|  | Mean | 12.95 | 13.16 | 12.74 |
|  | Std. Deviation | 2.97 | 3.02 | 2.92 |
| | Group Difference | $t(123) = 0.80, p = .423$ | | |
| Sex (N) | Male | 68 | 33 | 35 |
|  | Female | 57 | 30 | 27 |
|  | Total | 125 | 63 | 62 |
| | Group Difference | $\chi^2(1, N = 125) = 0.08, p = .782$ | | |
| Total brain volume (mm) | Range | 883847-1731175 | 912468-1452518 | 883847-1731175 |
|  | Mean | 1206619 | 1205468 | 1207789 |
|  | Std. Deviation | 124060.4 | 114255.4 | 134221.9 |
| | Group Difference | $t(123) = -0.14, p = .917$ | | |

To establish EAL as a metric of signal latency, we built models predicting N170 latency to both faces and symbols from EAL for each pathway (V1-rFG, V1-rpSTS). As sex(*41*), age(*42*), and brain volume(*43*) are known to impact brain structure and signal timing, we included these variables and their interactions with EAL as predictors in each model. Given the importance of the V1-rFG and V1-rpSTS pathways for face processing, we anticipated positive associations between N170 latency and EAL within these pathways during face but not symbol perception. We hypothesized that the association between N170 face latency and EAL along these pathways would be strongest for the typically developing control (TDC) group, indicative of a more efficient and effective face processing system. Because N170 latency is known to decrease throughout development(*20*), we anticipated a negative association between age and N170 latency for all pathways, conditions, and cohorts. To identify the model best suited for predicting N170 latency for each condition (face, symbol), pathway (V1-rFG, V1-rpSTS) and cohort (total sample, TDC, ASD), we performed stepwise model selection using both forward and backward regression, which involves the addition and removal of predictors until the model with the lowest Akaike Information Criterion (AIC) is found(*44*).

### Estimated axonal latency is a face-specific predictor of N170 latency

We first investigated associations between EAL and N170 latency along the V1-rFG and V1-rpSTS pathways for the total sample (Table 2, Fig. 2). For both pathways, the best fit models predicting N170 latency to faces revealed a significant positive association with EAL, and a significant negative association with age. These associations between N170 latency and EAL along both investigated face processing pathways were specific to faces. When predicting N170 latency to symbols, for both pathways, age was a significant negative predictor in the best fit models. While EAL remained a predictor for only the V1-rFG pathway, its association with N170 latency to symbols was not significant. In the total sample, brain volume and sex were never retained in any models for either pathway or condition.

**Fig. 2:**
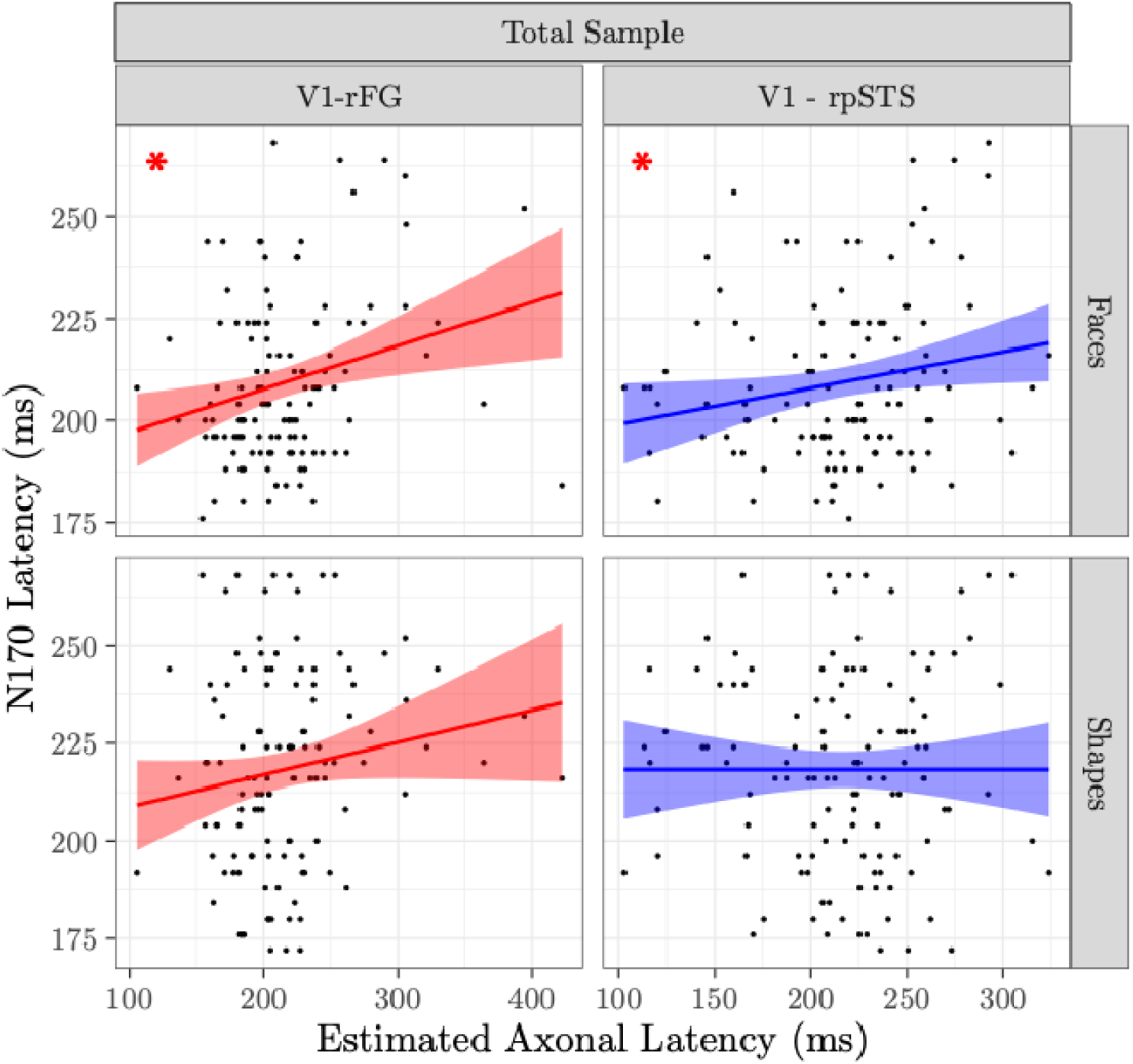
Associations between estimated axonal latency and N170 latency are specific to face processing along relevant pathways. The relationship between estimated axonal latency (EAL) and N170 latency is plotted for each pathway (red, V1-rFG; blue, V1-rpSTS) and each condition (faces, shapes) for the total sample (*N*=125). Results reveal a significant positive association between EAL and N170 latency across both pathways only in the faces condition. Significant associations are indicated via red asterisk. V1, primary visual cortex; rFG, right fusiform gyrus; rpSTS, right posterior superior temporal sulcus.

**Table 2:** Results of the best fit models predicting N170 latency for the total sample. A stepwise regression analysis revealed the best fit model for each condition and pathway in the total cohort by identifying the model with the lowest AIC value. Significant predictors are indicated via boldface font. Predictors that were not retained in the best fit model are indicated via hyphen.

| Condition | Pathway | Adjusted R<sup>2</sup> | Estimated Axonal Latency | Age | Sex | Brain Volume |
| --- | --- | --- | --- | --- | --- | --- |
| Faces | V1 - rFG | .21 | **b = 0.08, p = .023** | **b = -0.24, p < .001** | - | - |
|  | V1 - rpSTS | .21 | **b = 0.08, p = .022** | **b = -0.24, p < .001** | - | - |
| Shapes | V1 - rFG | .04 | b = 0.07, p = .162 | **b = -0.13, p = .043** | - | - |
|  | V1 - rpSTS | .04 | - | **b = -0.16, p = .015** | - | - |

### Individuals with and without autism rely upon different pathways for face processing

We next investigated associations between EAL and N170 latency along the V1-rFG and V1-rpSTS pathways for each cohort separately (Table 3, Fig. 3). When predicting N170 latency to faces, we found a significant positive association with EAL in the TDC cohort for the V1-rFG pathway, but EAL was not retained for the V1-rpSTS pathway. However, we found divergent results in the ASD cohort. When predicting N170 latency to faces for the ASD cohort, EAL within the V1-rFG pathway was not retained in the best fit model, but we found a trending positive association with EAL within the V1-rpSTS pathway. Age was a significant negative predictor in all best fit models of N170 latency to faces for both cohorts and pathways. For the ASD cohort only, sex remained an important, albeit non-significant, predictor for both pathways in these face latency models.

**Fig. 3:**
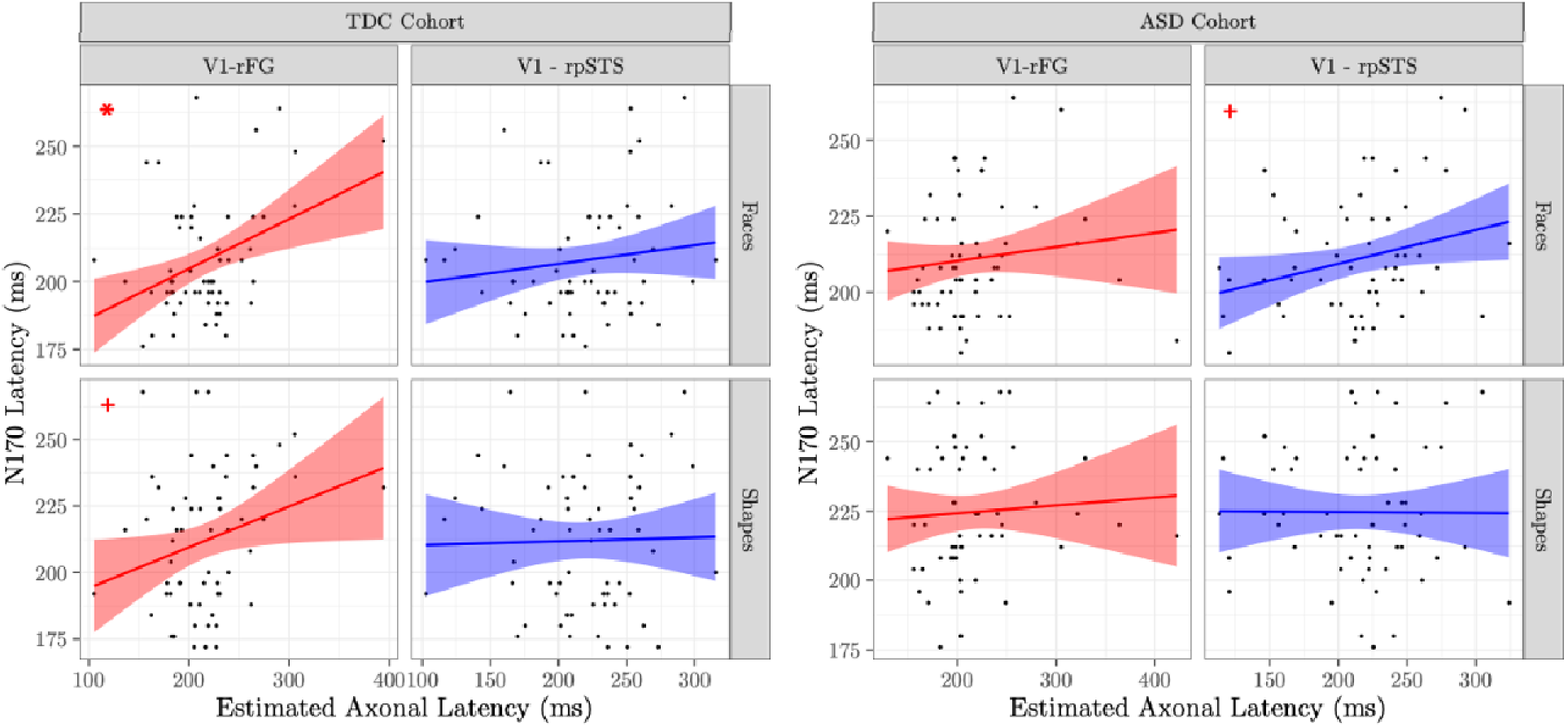
Estimated axonal latency is positively associated with N170 latency in different pathways for individuals with and without autism. The relationship between estimated axonal latency (EAL) and N170 latency is plotted for each pathway and each condition (faces, shapes) for the TDC cohort (*N*=63, left) and the ASD cohort (*N=*62, right). For TDC participants, we find EAL is positively associated with N170 latency to faces along only the V1-rFG pathway (red). For ASD participants, EAL is trending towards a positive association with N170 latency to faces along only the V1-rpSTS pathway (blue). Significant associations are indicated via red asterisk. Trending associations are indicated via red +. TDC, typically developing control; ASD, autism spectrum disorder; V1, primary visual cortex; rFG, right fusiform gyrus; rpSTS, right posterior superior temporal sulcus.

**Table 3:** Results of the best fit models predicting N170 latency for the TDC and ASD cohorts. A stepwise regression analysis revealed the best fit model for each condition and pathway in the TDC and ASD cohorts by identifying the model with the lowest AIC value. Significant predictors are indicated via boldface font. Trending significant predictors are indicated via italicized font. Predictors that were not retained in the best fit model are indicated via hyphen.

| Condition | Cohort | Pathway | Adjusted R<sup>2</sup> | Estimate<br>d Axonal<br>Latency | Age | Sex | Brain<br>Volume |
| --- | --- | --- | --- | --- | --- | --- | --- |
| Faces | TDC | V1 - rFG | .29 | **b = 0.15,**<br>**p < .001** | **b = -**<br>**0.25, p <**<br>**.001** | - | - |
|  |  | V1 -<br>rpSTS | .21 | - | **b = -**<br>**0.27,**<br>**p < .001** | - | - |
|  | ASD | V1 - rFG | .17 | - | **b = -**<br>**0.22,**<br>**p = .001** | b = 7.61,<br>p = .105 | - |
|  |  | V1 -<br>rpSTS | .20 | *b = 0.09,*<br>*p = .065* | **b = -**<br>**0.20,**<br>**p < .001** | b = 6.5,<br>p = .140 | - |
| Shapes | TDC | V1 - rFG | .08 | *b = 0.13,*<br>*p = .073* | b = -<br>0.14, p =<br>.130 | - | - |
|  |  | V1 -<br>rpSTS | .04 | - | *b = -*<br>*0.17,*<br>*p = .053* | - | - |
|  | ASD | V1 - rFG | .08 | - | b = -<br>0.14,<br>p = .130 | - | - |
|  |  | V1 -<br>rpSTS | .01 | - | b = -<br>0.11,<br>p = .211 | - | - |

No variables were significant predictors of N170 latency to symbols in either cohort or pathway. While not significant, EAL was retained in the model predicting N170 latency to symbols as a trending positive association for only the TDC cohort only along the V1-rFG pathway. Age was retained as a non-significant but negative predictor of N170 latency to symbols for both cohorts and pathways. Brain volume and sex were never retained in any models predicting N170 latency to symbols.

## Discussion

Neuroimaging modalities have traditionally captured proxies of brain function, such as changes in cortical blood volume or blood oxygen level dependent (BOLD) response. These measures are reflective of indirect metabolic and neurovascular demands that co-occur with functional activation at relatively low temporal resolution.(*45*) On the other hand, direct measures of neuronal electrical function, such as EEG, typically lack detailed spatial resolution and cannot examine the function of deep subcortical structures critical for complex behavior and implicated in disease processes. Historically, this has made it challenging to study the underlying neurophysiological correlates of disrupted cognitive function in brain disorders like ASD. For the first time, we present and validate a unified approach to estimate *in vivo* functional action potential transmission time between regions of the human brain using non-invasive, widely available, and spatially specific MRI techniques. This approach, estimated axonal latency (EAL), utilizes classic neuroscientific models of axonal structure, particularly diameter and myelin thickness, and traditional tractography methods to provide a voxel-wise estimate of action potential transmission time between regions of the brain. To validate this measure, we evaluated the association between EAL along two well established face processing pathways and face processing speed measured via EEG. We observed specific and significant relationships between N170 latency, a well-established electrophysiological metric of face processing speed, and EAL along the V1-rFG and V1-rpSTS pathways. By using adolescent populations with and without autism, we demonstrated its ability to account for differences in neuronal processing speed that occur across development and vary with disorder.

In a large, well characterized adolescent sample of both autistic and typically developing controls, EAL was specifically associated with face processing speed as measured by N170 signal latency– an association maintained across both investigated face processing pathways. In both V1-rFG and V1-rpSTS pathways, we find significant positive associations between N170 latency to faces and EAL. These results were nearly statistically identical in both the V1-rFG and V1-rpSTS pathways in the total sample, underscoring their joint importance for face processing and role in N170 signal generation.(*34*, *35*) Interestingly, the association between N170 latency and EAL was completely absent in response to non-face stimuli, demonstrating the noteworthy specificity of EAL to the investigated cognitive process along relevant pathways. The lack of a structure-function relationship during the non-specialized cognitive condition (i.e., symbol processing) may provide a structural account for the conflict resolution that must occur when the brain is deciding along which pathway to propagate a given signal when categorizing competing representations of incoming stimuli.(*46*)

We also found that the best fit models for each condition and pathway included age as a significant negative predictor of N170 latency in the total sample. This result is not surprising, as myelination of the cortex continues throughout adolescent development.(*42*, *47*) These developmental processes increase electrophysiological signaling speed and network efficiency.(*43*, *48*) However, despite the strong effect of age, EAL remained a significant positive predictor capable of explaining variance in N170 latency to faces above and beyond that explained by age.

We next examined EAL in the context of ASD, as autistic individuals often display slower N170 latencies to faces than their TDC counterparts(*22*) and demonstrate white matter differences in development.(*40*, *49*, *50*) To investigate whether our novel structural metric of neuronal latency may account for these observed functional differences, we investigated the relationship between EAL and N170 latency in the TDC and ASD cohorts separately. For N170 latency to faces, we observed divergent results between the two groups. For the TDC cohort, we found positive associations between N170 face latency and EAL along the V1-rFG pathway. However, for ASD participants, we find only a trending positive association between EAL and N170 face latency in the V1-rpSTS pathway. While FG and pSTS are both core nodes(*51*) in the face processing network that enable the successful perception, interpretation, and response to faces, these regions each contribute to unique aspects of face processing.(*35*) The V1-rFG pathway is the canonical, shorter face processing pathway recruited for the holistic processing of human faces. Alternatively, the V1-rpSTS is a longer and perhaps less direct pathway important for the processing of facial features fundamental to expression and identity.(*35*) While others have found similar relationships between fMRI BOLD activation to faces and N170 latency in both autistic and neurotypical individuals(*21*), our results demonstrate that V1-rFG EAL is a structural determinant of neural signal latency during face processing for TDC individuals only. Consequently, ASD participants may instead rely upon a more diffuse structural network for face processing, as V1-rpSTS EAL only weakly accounts for N170 latency in this cohort. These results provide a potential structural explanation of N170 latency delays often observed in this population.

Given our desire to account for differences in neuronal processing speed that occur in development and vary in autism, we included both sex and brain size as predictors in our models due to their documented associations with ASD(*52*, *53*) and development.(*42*) ASD diagnosis(*54*) and symptom presentation(*55*) have been shown to vary across sexes. For example, while females are often better able to camouflage or mask their symptoms,(*52*, *56*) differences in N170 face latencies have been associated with symptom severity in females, but not males.(*57*) Interestingly, sex was retained in models predicting N170 latency only to faces and only in the ASD cohort. This result indicates that sex is required to account for variance in face processing speed only for ASD individuals. Despite the fact that brain size increases reliably with age(*42*) and that brain volume differences are consistently observed in ASD,(*53*, *58*) total brain volume was not retained in any model included in our analyses. Therefore, brain size fails to account for additional variance in N170 latency above and beyond our novel EAL metric.

### Limitations and future directions

In calculating EAL, we used the T1-weighted/T2-weighted ratio, a widely available and easily acquired MRI metric to measure myelin content. However, there are limitations in the specificity of this metric for the quantification of myelin compared to gold standard magnetization transfer myelin quantification, particularly within deep white matter pathways.(*59*) However, with the significant associations between structural and functional metrics of signal latency identified in this study, we would only expect improved myelin quantification in future studies to reduce noise and improve EAL estimation. Future studies should expand upon our foundational work to include the investigation of additional pathways, cognitive processes, developmental stages, and disorders.

Another limitation to the current work is that we performed a secondary analysis of an EEG paradigm that presents emotional faces and symbols as feedback in the context of a visual implicit learning task.(*38*) These stimuli were presented concurrently with task feedback text (i.e., “Right” or “Wrong”), and they differed on low level stimulus features such as spatial frequency and contrast. As such, this paradigm was not optimized to assess basic face processing, and N170 latency may have been impacted by additional ongoing cognitive processes related to the central task or processing of the co-occurring visual information. Future work can address this limitation through use of a more optimal face processing paradigm.

Finally, while it is a strength that we employed a multimodal imaging approach to validate our novel EAL metric, future work can further interrogate associations between the structural and electrophysiological determinants of signal speed with dual EEG-fMRI data collection. We employed a powerful meta-analytic approach to define our V1, FG, and pSTS regions of interest, but results may be further strengthened by simultaneously capturing N170 latency and defining functional regions of interest on an individual level via fMRI.

Establishing the relationship between brain structure and function has been a key, decades-old goal in cognitive neuroscience as individual differences in structural and functional networks may shape the phenotypes of neurodevelopmental disorders.(*60*–*62*) Here, we introduce EAL as a novel metric capable of quantifying the structural determinants of neuronal signal latency. For the first time, we present a structural measure of neuronal function validated by electrophysiological measurements. By combining voxel-wise estimates of neuronal conduction velocity with measures of tract length, EAL is able to provide estimates of signal latency between any two connected regions in the brain at either the single fiber or whole-brain network level. Additionally, EAL can be derived on a single-subject level, paving the way for precision imaging estimates of individual processing time for specific cognitive functions. This approach not only allows for a deeper understanding of the normative relationship between processing speed and cognitive function, but also the ability to individually measure differences in this relationship which are often observed in heterogeneous neurodevelopmental disorders like ASD. Such advancements could pave the way for predictive biomarkers of atypical neural development and inform the mechanisms underlying targeted interventions aimed at improving cognitive and social outcomes for autistic individuals. By applying this metric to known cognitive networks, EAL could advance computational simulations of the brain derived from human data by providing accurate estimates for signal transmission time. As such, EAL provides an avenue towards significant advancement in human neuroimaging as a novel *in vivo* cellular approach to understanding complex human behavior.

## Materials and methods

### Participants

Here, we employ a large dataset collected at Seattle Children’s Research Institute, the University of California Los Angeles, Harvard Medical School, and Yale University as part of Wave 1 of the Gender Exploration of Neurogenetics and Development to Advance Autism Research (GENDAAR) consortium, supported by an NIH Autism Center of Excellence Network from 2013 to 2017. Institutional Review Boards at each site approved study procedures and participants and their parents provided informed consent/assent. Exclusion criteria for this dataset included premature birth, known genetic conditions, history of neurological disorder except uncomplicated non-focal epilepsy, active seizures within the last year, and an inability to comprehend task instructions. All participants underwent a comprehensive phenotyping process including the collection of demographic data such as sex and age as well as confirmation of autism diagnosis for the ASD cohort. The ASD cohort’s autism diagnosis was confirmed by a trained and research reliable clinician using both the Autism Diagnostic Interview, Revised (ADI-R)(*63*) and the Autism Diagnostic Observation Schedule, Second Edition (ADOS-2)(*64*). The TDC cohort was included in the dataset if they had no first-or second-degree relatives with autism, no developmental, neurological, or psychiatric disorders, and no evidence of elevated autism traits based on a total t-score <65 on a parent-report version of the Social Responsiveness Scale, Second Edition (SRS-2(*65*)).

ASD and TDC participants were included in the present study if they had completed the EEG implicit learning task and underwent structural and diffusion MRI (*n*=198). After completing the data preprocessing steps outlined below, 125 participants (*n*=62 autistic) remained in the analysis with sufficient data across all required imaging modalities. Participant characteristics for this sample are detailed in Table 1.

## EEG methods

### EEG data collection

At all four sites, EEG data were collected with the Electrical Geodesics, Inc (EGI) 128-channel Net Amps 300 system (Magstim EGI Inc., Eugene OR) with HydroCel nets using Net Station 4.4.2, 4.5.1, or 4.5.2 with a standard Net Station acquisition template. Data were sampled at 500 Hz, referenced to Cz, and impedances were kept below 50 kΩ. During their visit, participants’ heads were measured and the appropriate size electrode net was applied before beginning EEG collection. Each participant was seated in front of an 80 cm monitor with a button box and researcher beside them. Participants underwent EEG during a series of paradigms including an implicit learning task(*38*) in which they were instructed to classify abstract, fractal images into two groups. Participants were provided feedback on their performance in the form of emotional faces from the NimStim face database(*66*) (120 trials) and nonsocial symbols (120 trials). These feedback stimuli were matched for visual angle only. Correct responses were indicated via a happy male or female face (social) or a colored star (nonsocial) presented in the center of the screen, and the word “Right” presented at the bottom of the screen. Incorrect responses were indicated via a sad male or female face (social) or a colored “X” (nonsocial) presented in the center of the screen, and the word “Wrong” presented at the bottom of the screen.

### EEG data processing

Data preprocessing was completed using the Automated Pipe-Line for the Estimation of Scale-wise Entropy from EEG Data (APPLESEED v1.2.20240717)(*39*) implemented with EEGLab v2022.1 and Matlab R2020a software (Mathworks, Natick, MA). In this pipeline, data were downsampled to 250 Hz and segmented into -100 to 1000 ms epochs time locked to feedback stimulus (face, symbol) onset. Problematic channels with excessive or flat amplitudes were removed, and then epochs were identified as artifacts and removed if (1) the experimenter indicated a trial was invalid (i.e., the participant was not attentive) during data collection, or (2) voltage exceeded 500 µV in any channel. The data were then subjected to independent component analysis, and problematic components were identified and removed using the adjusted_ADJUST algorithm.(*67*) On average, 24.96 components (4-62) were removed. Participants with greater than 45 components removed were excluded from further analysis (*n*=12, 6 TDC). Epochs in which the standard deviation exceeded 80 µV within a 200 ms sliding window with a 100 ms window step were then discarded as artifacts. Participants with fewer than 50 trials per condition were excluded from further analysis (*n*=22, 9 TDC). On average, 102.12 face trials (51-120) and 101.80 symbol trials (50-120) were retained. Finally, problematic channels identified via the FASTER algorithm(*68*) and those initially discarded as problematic were interpolated. On average, 6.08 channels (2-11) were interpolated.

### N170 latency computation

We quantified N170 latency to faces and symbols as the time at which the maximum negative peak over 3 consecutive points occurred 170 to 270 ms after stimulus onset in parietal/occipital channels over the right hemisphere (channels 89, 90, 91, 94, 95, 96).(*23*, *69*–*71*) Visual manual inspection confirmed successful peak identification for each participant and condition. For the total sample, N170 latency to faces averaged 209.80 (176-268) ms, and N170 latency to symbols averaged 217.26 (172-268) ms. For the TDC cohort, N170 latency to faces averaged 206.91 (176-268) ms, and N170 latency to symbols averaged 210.67 (172-268) ms. For the ASD cohort, N170 latency to faces averaged 212.40 (180-268) ms, and N170 latency to symbols averaged 223.20 (176-268) ms. To investigate differences in N170 latency, we performed a repeated-measures ANOVA with the within-subjects factor of condition (faces, symbols) and the between-subjects factor of cohort (TDC, ASD) for the final dataset. As anticipated, this analysis revealed a significant main effect of cohort, such that TDC participants had significantly faster latencies than ASD participants (*F*(1,123)=6.47, *p*=.012), and a significant main effect of condition such that latencies to faces were significantly faster than latencies to symbols (*F*(1,123)=15.44, *p*<.001). There was also a significant interaction between cohort and condition (*F*(1,123)=4.72, *p*=.032) such that ASD participants displayed particularly slower latencies to symbols. Although the grand average N170 amplitude negativity was not greater for faces with this paradigm (Fig. 1A), only latency was of interest for the present analyses.

## MRI methods

### MRI collection

At all four sites, structural and diffusion images were acquired via a harmonized imaging protocol using either Siemens TrioTim or Prisma scanners with field strength of 3T. Diffusion images were acquired with an isotropic voxel size of 2×2×2mm^3^, 64 non-collinear gradient directions at b = 1000 s/mm^2^, and 1 b = 0, TR = 7300ms, TE = 74ms. T1-weighted MPRAGE images were acquired with a FOV of 176×256×256, an isotropic voxel size of 1×1×1mm^3^, and TE = 3.3. T2-weighted images were acquired with a FOV of 128×128×34 with a voxel size of 1.5×1.5×4mm^3^, and TE = 35.

### MRI image preprocessing

T1-weighted and T2-weighted images were corrected for bias fields using the N4 algorithm implemented in ANTs.(*72*) T1-weighted images were processed using the *recon-all* pipeline in Freesurfer(*73*) in order to generate volume measurements and to generate individualized brain masks. Diffusion images were preprocessed according to established protocols that have been shown to be highly reliable means of estimating white matter fiber orientation and microstructural information.(*74*) Images were denoised,(*75*) corrected for Gibbs rings,(*76*) and processed for motion correction using the *eddy* command in FSL(*77*) incorporating nonparametric outlier detection and replacement. Subjects were then visually checked for quality control and individuals with large errors excluded from further analysis (*n*=4, 1 TDC). Diffusion images were upsampled to 1.3×1.3×1.3mm isotropic voxel size to match Human Connectome Project data standards, and response functions for white matter, gray matter, and cerebrospinal fluid tissue types were generated using the *Dhollander* algorithm implemented in MRtrix3 from a subgroup of 40 randomly selected age-matched participants counter-balanced for sex and diagnosis. The fiber orientation distribution (WM-FOD) was calculated via constrained spherical deconvolution using the response functions and the MRtrix3Tissue package (https://3tissue.github.io).

### Region of interest selection and creation

We defined our V1, rFG, and rpSTS regions of interest (ROIs, Fig. 1C) with a meta-analytic approach using Neurosynth.org.(*78*) V1, rFG, and rpSTS were chosen because of their well-established role in the facial processing network and visual processing stream.(*34*, *35*) V1 was extracted from the Neurosynth term-based meta-analysis for “primary visual,” thresholded at a z-score ≥= 3.5, and binarized. FG was extracted from the term-based meta-analysis for “ffa,” thresholded at a z-score ≥= 12.0, binarized, and cropped to isolate the right hemisphere. pSTS was extracted from the term-based meta-analysis for “psts,” thresholded at a z-score ≥= 3.5, binarized, and cropped to isolate the right hemisphere. All masks were registered to align with the T1-weighted image for each subject, and shifted anteriorly by 12 mm via linear transform to avoid overlapping the cerebellum. All cropping and thresholding was done using FSL’s FSLview image editing tools and FSLmaths.(*77*)

### Tractography

Whole brain tractography was performed using MRtrix3 similar to previously published methods.(*79*) The WM-FOD was used to provide directional information for fiber tracking and density. The white matter/gray matter interface was determined for each subject derived from their T1-weighted image using a hybrid surface and volume approach including subcortical structures.(*80*) The interface was seeded with streamlines until 50,000,000 tracts successfully meeting inclusion criteria were generated. These were subsequently pruned back to 10,000,000 tracts via spherically informed filtering of tractograms (SIFT), a method to preserve anatomical fidelity by matching the density of tracts traversing each voxel to the WM-FOD lobe integral.(*81*) The WM-FODs were used to register each subject to the group-wise template, then to stereotaxic space.(*82*) The resulting warps were used to move the ROIs into subject space. Tracts were retained if the coordinates of each endpoint terminated within the ROI of interest (rFG, rpSTS) on one end and V1 on the other. Exclusion criteria removed tracts that were located in the cerebellum or crossed the corpus callosum. The overall mean length of the retained tracts was calculated via the *tckstats* command implemented in MRtrix3, which sums the distance between each component vertex along the length of each retained tract. This was performed in native space to avoid any errors arising from altering vertex distance after warping images to a common space. Participants were excluded if no tracts were retained between the ROIs. Thirty-nine participants (20 TDC) were excluded due to no tracts remaining along the V1-rpSTS pathway, and an additional 10 participants (5 TDC) were also excluded from V1-rFG analyses due to no tracts remaining along this pathway. To investigate differences in tract length, we performed a repeated-measures ANOVA with the within-subjects factor of pathway (V1-rFG, V1-rpSTS) and the between-subjects factor of cohort (TDC, ASD). This analysis revealed a significant main effect of pathway, such that the V1-rpSTS pathway was significantly longer than the V1-rFG pathway (*F*(1,117)=20.95, *p*<.001). There was no significant effect of cohort (*F*(1,117)=1.38, *p*=.243), and no significant interaction between pathway and cohort (*F*(1,117)=0.16, *p*=.686). Voxels traversed by the retained tracts were identified and the aggregate conduction velocity within these voxels was averaged for each subject.

### Estimated axonal latency (EAL) calculation

EAL is derived from G-ratio and aggregate conduction velocity metrics. Both metrics were calculated according to previously published methods(*40*). The aggregate g-ratio was calculated on a voxel-wise basis according to Stikov et al.(*83*, *84*) and was used according to Mohammadi & Callaghan,(*13*) as displayed in Equation 1. As a measure of intra-axonal volume, the fiber density cross section was used as the axonal volume fraction (AVF).(*85*) As a metric of myelin density, the T1-weighted/T2-weighted ratio was used as the myelin volume fraction (MVF). These metrics represent the total sums of each respective compartment across the volume of the voxel and are a volume-based equivalent to the original formulation of g as the ratio of axon diameter (d) to fiber diameter (D).

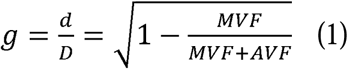

Aggregate conduction velocity (ACV) was calculated based on the calculations of Rushton(*86*) and Berman et al.,(*14*) reiterating Rushton’s calculation that conduction velocity (θ) is proportional to the length of each fiber segment (l) and that this is roughly proportional to *D,* which in turn can be defined as the ratio between *d* and the g-ratio (g). Furthering the considerations of Rushton, Berman et al. show that a value proportional to conduction velocity can be calculated using axon diameter and the g-ratio as in equation 2:(*14*)

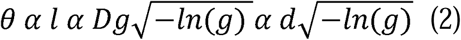

To investigate differences in aggregate conduction velocity, we performed a repeated-measures ANOVA with the within-subjects factor of pathway (V1-rFG, V1-rpSTS) and the between-subjects factor of cohort (TDC, ASD). This analysis revealed a significant main effect of pathway, such that aggregate conduction velocity was significantly faster along the V1-rpSTS pathway (*F*(1,117)=39.96, *p*<.001). There was no significant effect of cohort (*F*(1,117)=0.81, *p*=.370), and no significant interaction between pathway and cohort (*F*(1,117)=0.03, *p*=.853).

Finally, EAL is calculated according to equation 3, which is derived from the equation for average velocity, a well-known relationship in classical physics.

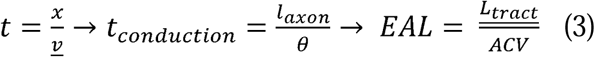

It has often been assumed that action potentials propagate down axons at roughly a constant speed(*2*). Thus, we can relate the conduction velocity of an action potential to the axonal length and conduction latency. The aggregate conduction velocity of a given tract is proportional to its true conduction velocity, allowing us to substitute axonal length with mean tract length and conduction latency with theoretical latency. As a result, EAL should be proportional to conduction latency along a given tract.

To investigate differences in EAL, we performed a repeated-measures ANOVA with the within-subjects factor of pathway (V1-rFG, V1-rpSTS) and the between-subjects factor of cohort (TDC, ASD). This analysis revealed no significant effect of pathway (*F*(1,117)<0.01, *p*=.958), no significant effect of cohort (*F*(1,117)=0.09, *p*=.769), and no significant interaction between pathway and cohort (*F*(1,117)=0.05, *p*=.831).

### Statistical analysis

In order to identify the model best suited for predicting N170 latency across cohorts (total sample, TDC, ASD), pathways (V1-rFG, V1-rpSTS), and conditions (faces, symbols), we performed stepwise model selection using both forward and backward regression, which involves the addition and removal of predictors until the model with the lowest AIC criterion is found. Model selection was conducted using R 4.2.3(*87*) with the stepAIC function from the MASS package.(*88*) Our models included seven predictors chosen for their known impact on brain structure and function:(*41*–*43*) EAL, sex, age, total brain volume, and the interactions between EAL and each of the other predictors. We identified the best fit model as the model with the lowest AIC value before an intercept-only model. Finally, a general linear model was run on the predictors remaining in the best fit model to determine the direction and significance of its association with N170 latency.

## Funding

NIH R01MH100028 (KAP)

NIH K01MH125173 (MHP)

## Author Contributions

All data were collected by the Gender Exploration of Neurogenetics and Development to Advance Autism Research (GENDAAR) consortium. CRC and MGN contributed equally as first authors. BTN and MHP contributed equally as senior authors.

Conceptualization: CRC, MGN, BTN, MHP

Formal analysis: CRC, MGN, BTN, MHP

Investigation: VA, AK, CAWS, MS, EN, HB, RAB, SYB, MD, AJ, NMK, SJ, JCM, AN, DG, ARG, SJW, KAP, JVH

Data curation: ZJ

Writing–original draft: CRC, MGN, BTN, MHP

Writing–review & editing: CRC, MGN, MD, AJ, NMK, JCM, AN, SJW, KAP, JVH, BTN, MHP

Supervision: TJD, JVH, KAP, BTN, MHP

Funding: KAP, MHP

## Data Availability

All data are available through the NIMH Data Archive (NDA), at https://nda.nih.gov/edit_collection.html?id=2021.

